# Signal peptidase complex mediates rotavirus VP7 processing and virion assembly

**DOI:** 10.1101/2025.11.04.686468

**Authors:** Xuejiao Zhu, Liliana Sanchez-Tacuba, Wandy Beatty, Bin Li, Siyuan Ding

## Abstract

For viruses that replicate in the proximity of or bud at the endoplasmic reticulum (ER) associated membranes, proper processing of their glycoproteins is critical for successful infection. Rotavirus outer capsid protein VP7 is an ER-resident protein. However, its N-terminal signal peptide is removed by an unknown proteolytic mechanism. In this study, we leveraged tandem affinity purification followed by high-resolution mass spectrometry to profile host proteins that interact with VP7. We identified members of the signal peptidase complex (SPC) family as important host factors that facilitate rotavirus infection. CRISPR knockout or siRNA knockdown of distinct SPC subunits resulted in significant decrease in infectious rotavirus titers in a viral strain- and cell type-independent manner. While viral transcription, translation, and replication were not altered in the absence of SPC, we observed formation of abnormal viral particles by transmission electron microscopy (TEM) in *SPCS1* knockout cells. Mechanistically, loss of SPC proteins led to inefficient cleavage of VP7 signal peptide and severely impaired the final steps of virion maturation and assembly. Additionally, we identified residue E256 within VP7 as a key site for SPC binding. An E to R mutation abolished VP7 interaction with SPC and subsequently led to reduced viral infectivity. Taken together, these findings define SPC as a novel regulator of VP7 maturation and rotavirus assembly and highlight its role as a novel cellular target for potentially broad-spectrum antiviral therapeutic development.

**Author summary:** Previous reports demonstrated that SPC is involved in the replication cycles of several members of the flavivirus family and human T-cell leukemia virus type 1 (HTLV-1). As an ER-resident outer capsid protein, rotavirus VP7 must undergo proper post-translational modifications, including the cleavage of its N-terminal signal peptide, to be functionally incorporated into mature virions. However, the processing mechanism remains unknown. For the first time, our study identifies SPC as an essential regulator of rotavirus assembly by mediating the cleavage of the VP7 signal peptide. Loss of SPC impairs VP7 signal peptide cleavage and maturation, thereby disrupting correct virion assembly and reducing infectious particle production. Using Alphafold3, we predicted the VP7 residue E256 to be at the binding interface with SPC complex. Experimentally, mutation of glutamic acid to arginine (E256R) substantially weakens this interaction and results in reduced viral propagation. Our findings unveil a novel post-translational checkpoint in rotavirus regulated by SPC and underscore the promise of SPC as a broad-spectrum antiviral target, especially for rotavirus, flavivirus, and HTLV-1, whose viral glycoproteins and structural proteins require SPC processing for proper maturation.

## Introduction

Signal peptidase complex (SPC) is responsible for the cleavage of signal peptides of many secreted and membrane-associated proteins. The eukaryotic SPC consists of five subunits, including three non-catalytic subunits, signal peptidase complex subunit 1 (SPCS1), SPCS2 and SPCS3, and two catalytic subunits, including either SEC11A or SEC11C [1]. Among the SPC subunits, SEC11 and SPCS3 are essential for signal peptidase activity and cell survival [1, 2]. SPCS1 is a crucial host factor involved in the endoplasmic reticulum (ER)-based processing of both cellular and viral proteins. Recent studies identified SPCS1 as a proviral host factor, particularly for certain enveloped viruses that rely on the host’s ER machinery for the maturation of their structural proteins. Several viruses produce glycoproteins or structural proteins that require cleavage of signal peptides in the ER. SPCS1 is essential for processing these viral proteins, enabling their correct folding and transport to the cell surface or assembly sites. Viruses in the *Flaviviridae* family, such as Dengue virus (DENV), Zika virus (ZIKV), West Nile virus (WNV), and Japanese Encephalitis virus (JEV) have been shown to depend on SPCS1 for the proper processing of their envelope (E) and precursor membrane (prM) proteins [3-5]. Knockdown of SPCS1 impairs viral replication by blocking the maturation of these glycoproteins. Likewise, SPCS1, SPCS2, and SPCS3 were identified as proviral host factors involved in the late step of Hepatitis C virus replication [6]. In a newly published report, p12 protein of human T-cell leukemia virus type 1 (HTLV-1) is cleaved by the SPC to generate p8. Inhibition of SPC resulted in impaired HTLV-1 transmission, implicating SPC as a new cellular target to counteract HTLV-1 cell-cell transmission [7].

Rotavirus is a non-enveloped, icosahedral particle composed of three concentric protein layers that surround the segmented, double-stranded (ds) RNA genome [8, 9]. The innermost layer known as single-layered particles (SLPs) is formed by VP2, which encapsidate the viral dsRNA genome, the RNA-dependent RNA polymerase VP1 and the RNA capping enzyme VP3. Surrounding this core is the intermediate layer composed of VP6, the most abundant structural protein, which confers group and subgroup antigenic specificity and forms the double-layered particles (DLPs). RNA replication and DLP formation take place in cytoplasmic inclusions known as viroplasms [9]. DLPs then bud into the ER, where the outermost glycoprotein VP7 and spike protein VP4 assemble and mature into triple-layered particles (TLPs), then released by cell lysis. VP7 provides structural stabilization of the mature virion, while VP4 mediates attachment, entry, and proteolytic activation required for infectivity [10-15]. This highly ordered TLP design is essential for the protection of the viral genome and the efficient initiation of a new round of infection.

The viral glycoprotein, VP7, is therefore an ER resident glycoprotein with a luminal orientation [16-18]. Domain I (residues 78-161, 256-312) features a Rossmann-like α/β fold, which is a common structural motif found in nucleotide-binding proteins. It forms the structural backbone of the VP7 monomer and provides stability to the trimeric assembly. Domain II (residues 162-255) is inserted into domain I and adopts a jelly-roll β-barrel fold and contributes to the outer surface topology of VP7, including loops that serve as neutralizing epitopes. N-terminal region (Signal Peptide, residues 1–50) is cleaved off in the ER during maturation [19], which is essential for ER targeting and correct processing of VP7. The site of cleavage has been pinpointed to between Ala50 and Gln51. A single point mutation that converts Ala50 to Val prevented VP7 processing [16]. However, the cellular and molecular mechanisms involved in this process and the host enzymes responsible have not been determined.

Here, by interactome analysis, we found that several SPC family members interact with VP7. Knocking out endogenous SPCS1 and knocking down SPC family members significantly reduce production of rotavirus progenies. SPCS1 is important for post-translational protein processing and virus assembly rather than transcription or translation. Together with knowledge of the roles of VP7 in TLP formation and virion assembly, our results demonstrate for the first time that the host factor SPCS1 participates in the virion assembly of rotaviruses through interactions with VP7. Understanding the role of SPC in rotavirus infection offers new insights into host-pathogen interactions and instructs the design of host-targeted antiviral strategies.

## Results

### SPC interacts with rotavirus VP7

A quantitative proteomics approach was used to construct a comprehensive interactome network of host factors that co-precipitate with VP7 derived from human rotavirus DS1 strain. We adopted a similar method of tandem affinity purification followed by liquid chromatography mass spectrometry (LC-MS/MS) that was previously used in the interactome analysis for rhesus rotavirus NSP1 [20] and VP4 [21]. Upon two rounds of immunoprecipitation, thirteen proteins were found to be high-confidence host factors that interact with VP7. Among the thirteen proteins, five host proteins, including SPCS1, SPCS2, SPCS3, SEC11A, and SEC11C, belong to the SPC family, indicating strong interaction of the SPC with VP7 protein (**Fig 1A**, **Table 1, and Dataset S1**). To narrow down the top hits for follow-up analysis, we conducted co-localization analysis for some of the newly identified proteins. SPCS2 and SEC11C both strongly colocalized with VP7 in the ER (**Fig 1B and S1 Fig**), to comparable levels as the positive controls NSP1 and SAMD9 [22], VP3 and PFDN4 [23], and much higher than VP6 and ECE1, which served as a negative control. To further verify the interaction between SPC and VP7, we co-transfected FLAG-tagged SPCS1 and GFP-tagged VP7 in Human embryo kidney HEK293 cells and performed immunoprecipitation (IP) and immunoblot. We found that rotavirus VP7 could interact with host SPCS1 (**Fig 1C**), further provided the evidence that SPC interacts with the rotavirus VP7 protein.

**Fig 1.**
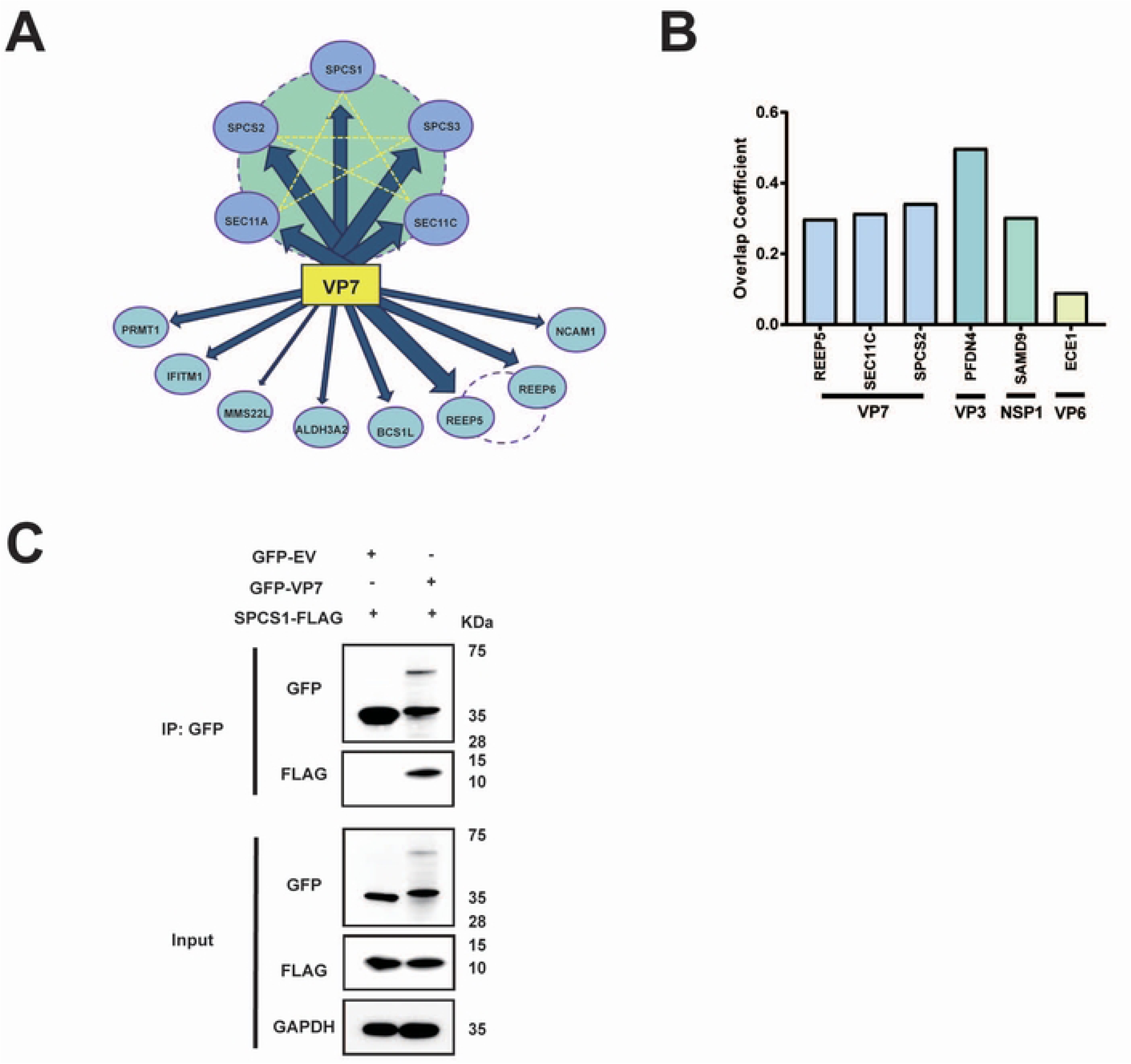
SPC interacts with rotavirus VP7. (A) Interactome of VP7 and host proteins. The proteins connected by dotted lines belong to the same family based on published interactome studies. The width of the arrows indicates the strength of the interaction between host proteins and DS1 VP7. (B) Co-localization analysis of VP7 and REEP5, SEC11C, SPCS2, VP3 and PFDN4, NSP1 and SAMD9, VP6 and ECE1. Results are shown as overlap coefficients based on images in Fig. S1. (C) Co-IP of SPCS1 with VP7. HEK293 cells were co-transfected with plasmids expressing FLAG-tagged SPCS1 and GFP-tagged DS1-VP7. Cell lysates were immunoprecipitated with an anti-GFP antibody. The resulting precipitates and whole-cell lysates used for immunoprecipitation were examined by immunoblot with anti-GFP and anti-FLAG antibodies.

**Table 1.** Peptide counts in VP7 interactome. This table summarizes the peptide counts of proteins identified in VP7 interactome. The target genes, their corresponding symbols, and the number of peptides detected for each are shown.

| Gene | Symbol | Count numbers |
| --- | --- | --- |
| signal peptidase complex subunit 2 | SPCS2 | 190 |
| signal peptidase complex subunit 3 | SPCS3 | 114 |
| signal peptidase complex catalytic subunit SEC11A isoform 3 | SEC11A | 70 |
| signal peptidase complex catalytic subunit SEC11C | SEC11C | 31 |
| signal peptidase complex subunit 1 | SPCS1 | 13 |
| receptor expression-enhancing protein 5 | REEP5 | 70 |
| receptor expression-enhancing protein 6 | REEP6 | 14 |
| protein MMS22-like isoform X6 | MMS22L | 29 |
| neural cell adhesion molecule 1 isoform 4 precursor | NCAM1 | 22 |
| fatty aldehyde dehydrogenase isoform 2 | ALDH3A2 | 20 |
| mitochondrial chaperone BCS1 | BCS1L | 18 |
| interferon-induced transmembrane protein 1 | IFITM1 | 6 |
| chromatin target of PRMT1 protein isoform X1 | PRMT1 | 4 |

### Loss of SPC impairs rotavirus growth

To investigate the role of endogenous SPCS1 in the propagation of rotavirus, we infected wild-type (WT) and *SPCS1* knock-out (KO) HEK293T cells with rhesus rotavirus (RRV) and measured virus yield. The viral titer in *SPCS1* KO cells was approximately 13-fold lower than that in WT cells and partially restored in the KO cells complemented with Flag-tagged SPCS1 (**Fig 2A and B**). Next, we evaluated whether the pro-viral effect of SPCS1 was rotavirus strain-dependent. Similar to the results obtained with RRV with 7.5-fold reduction in titers, infection with simian strain SA11 or bovine strain UK also showed 11-fold and 9-fold reduction viral titers in *SPCS1* KO cells compared with WT cells, respectively (**Fig 2C**). We validated these findings independently in Huh7.5 hepatoma cells. Consistently, UK, RRV, and SA11 strains showed 11.9-fold, 8.2-fold and 7.8-fold reduction viral titers in *SPCS1* KO Huh7.5 cells, respectively, compared with those in WT cells (**Fig 2D**), demonstrating that SPCS1 deficiency broadly impairs the propagation in a viral strain independent manner, the impairment of rotavirus propagation by the loss of SPCS1 was specific. To verify whether other members in SPCS family exert similar effects on RV propagation, siRNAs targeting *SEC11A*, *SEC11C*, *SPCS2* were transfected in HEK293T *SPCS1* KO cells and WT cells, achieving approximately 90% knockdown efficiency for SEC11A and SEC11C **(S2A-B Fig)**, and 80% for SPCS2 **(S2C Fig)**. We found that virus titers in *SPCS1* KO HEK293T cells were further reduced by 3.2-, 5-, and 6-fold following knockdown of *SEC11A*, *SEC11C*, and *SPCS2*, respectively (**Fig 2E**), suggesting that SPC complex had some residual activity without the non-catalytic subunit SPCS1. Taken together, it demonstrated that the SPC loss could impair rotavirus propagation.

**Fig 2.**
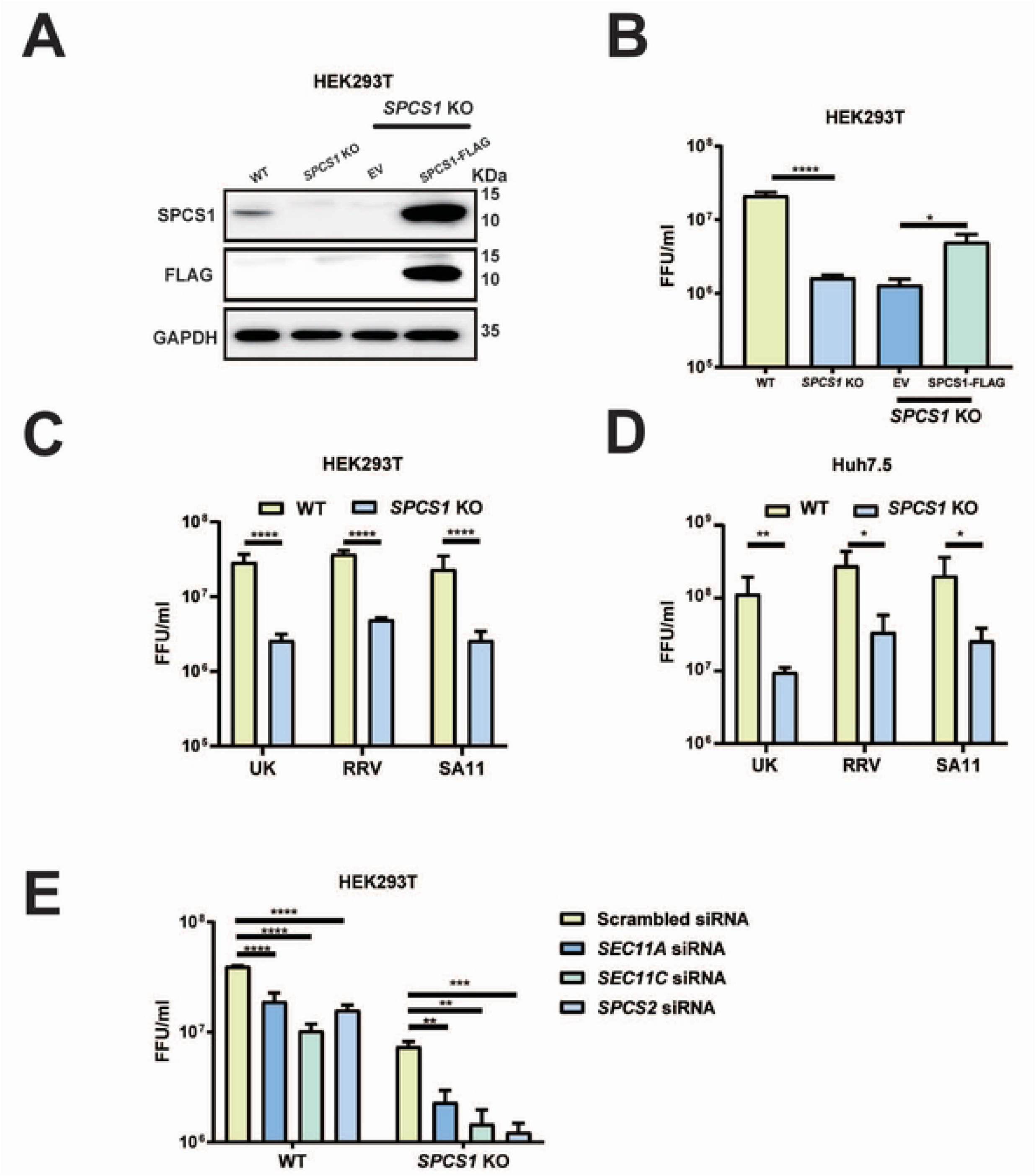
Loss of SPC impairs rotavirus growth. (A) WT, *SPCS1* KO HEK293T, *SPCS1* KO cells transfected with empty vector or FLAG-tagged SPCS1 were infected with RRV at an MOI of 3, and cells were collected for western blot analysis at 12 hpi using anti-FLAG and SPCS1 antibodies to confirm the protein level of SPCS1. (B) The lysates and supernatants were collected for FFU assays to determine the titers. (C) WT and *SPCS1* KO HEK293T cells were infected with RRV, SA11 and UK strains at an MOI of 3. At 12 hpi, all the cell lysates and supernatants were collected for FFU assay to detect the titers. (D) WT and *SPCS1* KO Huh7.5 cells were infected with RRV, SA11 and UK strains at an MOI of 3, respectively. At 12 hpi, all the cell lysates and supernatants were collected for FFU assay to detect the titers. (E) WT and *SPCS1* KO HEK293T cells were transfected with siRNAs targeted against *SPCS2*, *SEC11A*, *SEC11C* and a scrambled siRNA at the concentration of 20 nM. At 48 hpi, cells were infected with RRV at an MOI of 3. The cell lysates and supernatants were collected for FFU assay to detect the titers at 12 hpi. The results are the averages of data in three independent experiments and plotted as mean ± SD. Statistical significance was determined using two-way ANOVA with Sidak’s multiple comparisons test (**, *P* < 0.01; ***, *P* < 0.001, **** *P* < 0.0001).

### SPCS1 does not regulate viral RNA and protein levels during rotavirus infection

To determine the stage of the viral replication cycle was affected by SPCS1, we assessed viral infectivity by measuring NSP5 mRNA and VP6 protein levels in WT and *SPCS1* KO HEK293T cells at 4, 8 and 12-hour post infection. We found no significant differences in intracellular viral RNA levels comparing WT and *SPCS1* KO HEK293T cells or Huh7.5 cells at any time point (**Fig 3A, S3A Fig**). We also found VP6 protein levels remained unchanged between WT and *SPCS1* KO HEK293T cells, as well as WT and *SPCS1* KO Huh 7.5 cells at 4, 8 and 12 hpi (**Fig 3B, S3B Fig**). To further investigate whether SPCS1 influences rotavirus replication in viroplasm, we performed immunofluorescence staining to detect NSP5, a key marker of viral replication level within viroplasms in WT and *SPCS1* KO cells. There was no significant difference in the numbers and sizes of viroplasms between WT and *SPCS1* KO cells (**Fig 3C**). Together, these findings suggest that SPC is not involved in viral transcription and translation or any earlier steps including virus entry.

**Fig 3.**
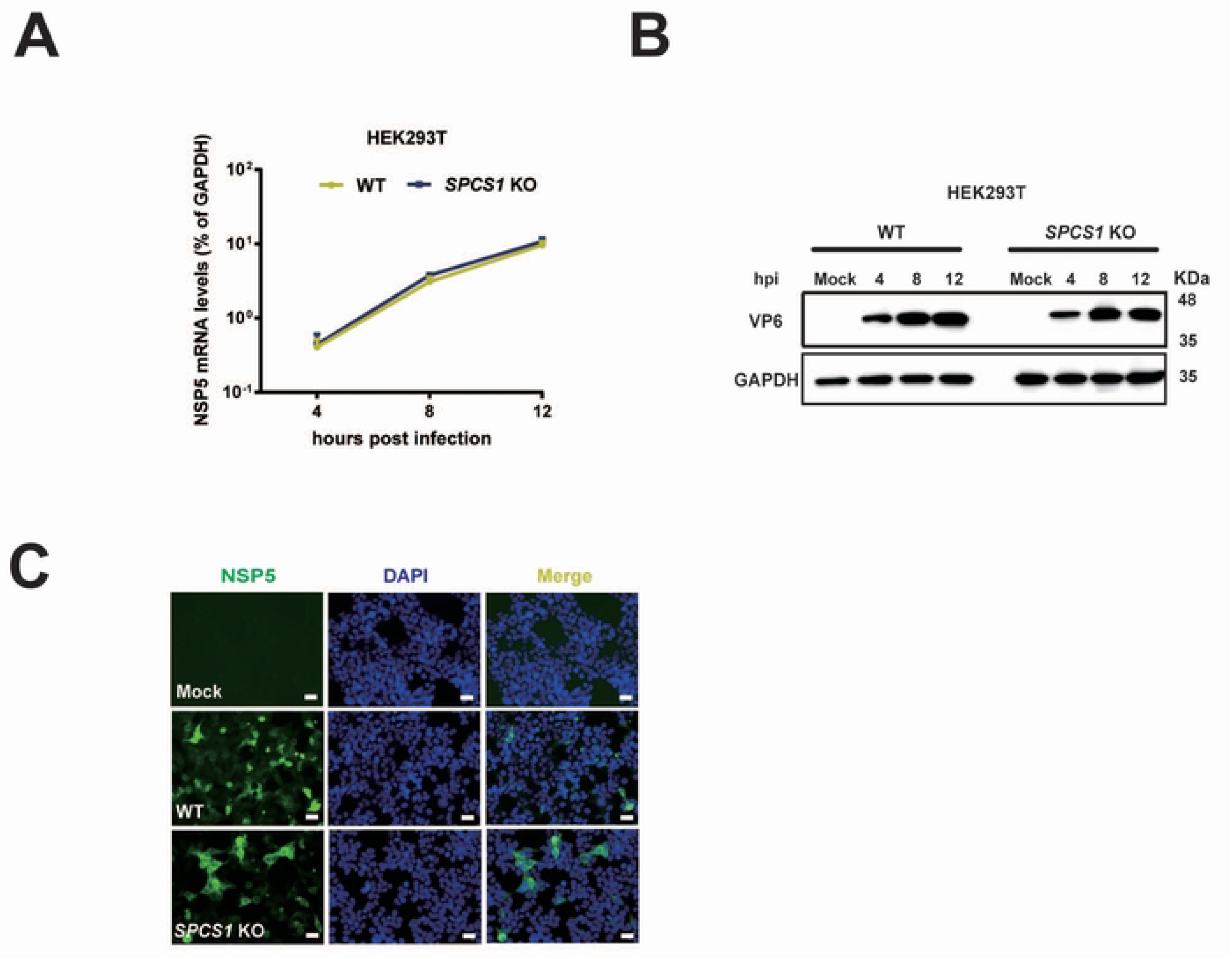
SPCS1 does not regulate RNA and protein level during rotavirus infection. (A) WT and *SPCS1* KO HEK293T cells were infected with RRV at an MOI of 3. At 4, 8, 12 hpi, the cells were collected for qRT-PCR analysis of viral mRNA level by detecting NSP5. The NSP5 level was normalized to that of GAPDH. Results are the average of data in triplicates and plotted as mean ± SD. (B) WT and *SPCS1* KO HEK293T cells were infected with RRV at an MOI of 3. At 4, 8, 12 hpi, the infected cells and mock cells were harvested for western blot analysis of viral protein level by detecting VP6 protein. (C) WT and *SPCS1* KO HEK293T cells were infected with RRV at an MOI of 3. At 12 hpi, an immunofluorescence assay was performed to detect the viroplasms in the infected cells. Cells were fixed and probed with in-house polyclonal antibody against NSP5; the positive cells were shown as green fluorescence. Nuclei were counterstained with DAPI. Representative data from one of three independent experiments are shown. Scale bar, 50 μm.

### Loss of SPCS1 hinders rotavirus TLP formation

Since the loss of SCPS1 did not affect rotavirus replication per se, we next examined later steps of viral replication cycle, in particular virus assembly. To that end, we purified the viral particles produced from RRV-infected WT and *SPCS1* KO HEK293T cells, and used the particles produced from MA104 cells as a positive control. The purified viral particles were subjected to SDS-PAGE and silver staining. We performed silver staining of viral proteins from purified rotavirus particles isolated from WT and *SPCS1* KO HEK293T cells, and MA104 cells. Major structural proteins, including VP1, VP2, VP3/VP4, VP6, and VP7, were clearly visualized. Compared to MA104 and WT HEK293T cells, viral particles from *SPCS1* KO cells exhibited reduced levels of outer capsid proteins VP4 and VP7 (**Fig 4A**). We confirmed this finding with western blot analysis of the purified viral particles. Consistent with the silver staining results, the structural protein VP4, VP7 and VP6 could be detected in MA104 cells when probed with an anti-TLP antibody. In contrast, VP7 signals seemed to be completely missing from *SPCS1* KO HEK293T cells (**Fig 4B**), indicating impaired expression or stability of TLPs with the loss of SPCS1. To definitively demonstrate that TLP formation was impaired, we infected WT and *SPCS1* KO cells with RRV and subjected them to transmission electron microscopy (TEM) analysis. We observed all forms of rotavirus particle formation, including SLPs and DLPs, TLPs and the budding process of DLP morphing into TLP (**S4A-D Fig**). In WT cells infected with rotavirus, we found a substantial number of viral particles close to the viroplasm (**Fig 4C**). Most of these particles were mature virions with a fully-formed triple layered structure (**Fig 4D**). Importantly, in *SPCS1* KO cells, although an equal number of particles surrounded the viroplasm (**Fig 4E**), with a similar size and number of viroplasms as observed with the previous immunofluorescence assay (**Fig 3C**), the morphology of these viral particles was notably aberrant, with the TLP structure severely disrupted, indicating a failure in proper viral assembly (**Fig 4F**). These results strongly suggest that SPCS1 loss leads to the abnormal TLP assembly and that SPCS1 is required for proper formation of TLPs during rotavirus assembly.

**Fig 4.**
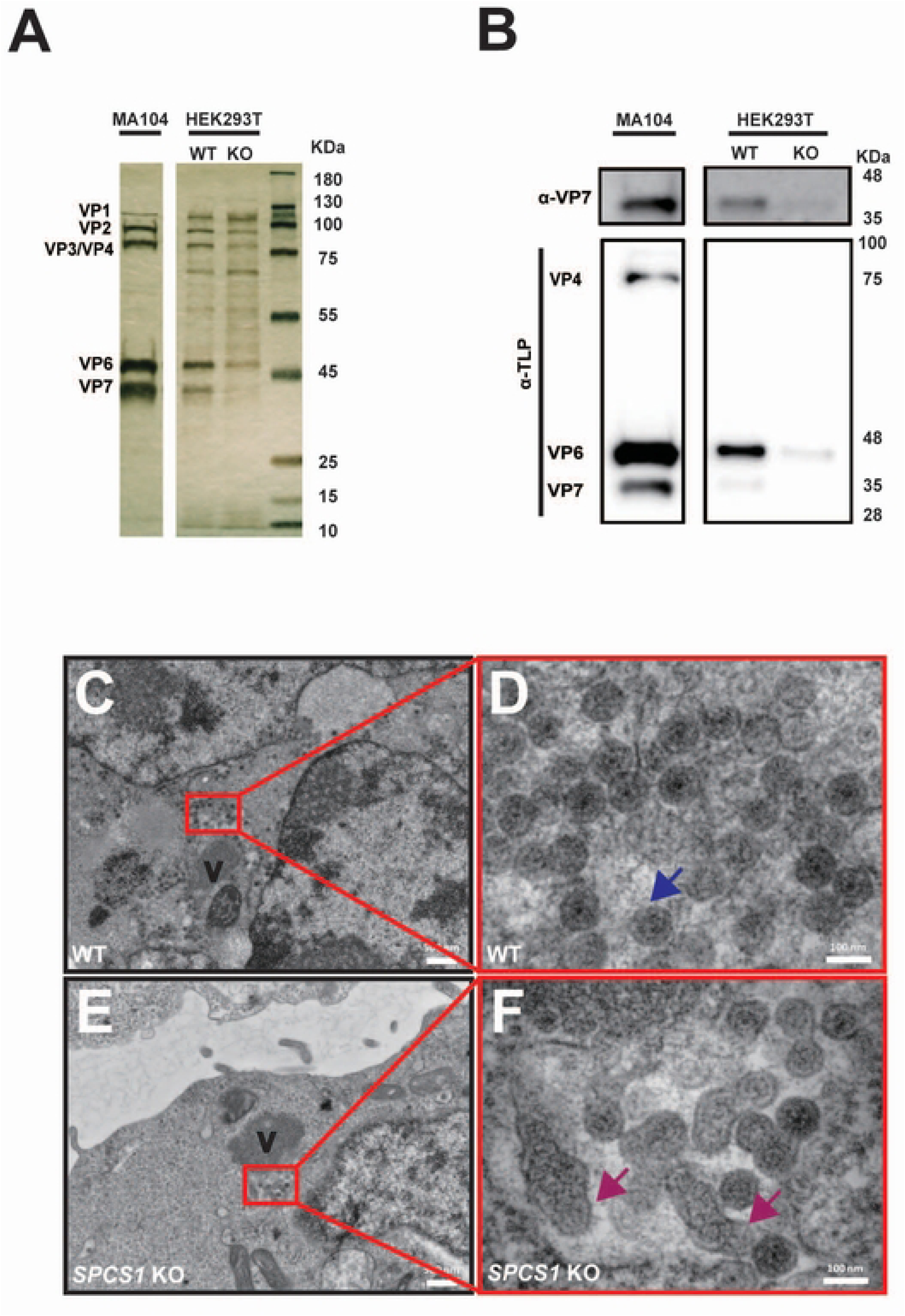
Loss of SPCS1 hinders rotavirus TLP formation. (A) WT and *SPCS1* KO HEK293T cells were infected with RRV at an MOI of 3 for 12 hours, and the supernatants were subjected to density gradient ultracentrifugation. The TLP layer was extracted following CsCl gradient ultracentrifugation and detected by SDS-PAGE and followed by silver staining. (B) Western blot validation of TLPs from WT and *SPCS1* KO HEK293T cells by probing TLP and VP7. (C) Electron micrograph (low magnification) of RRV-infected HEK293T WT cells. ‘V’ indicates the viroplasm. (D) Enlarged view of the area boxed in red in panel (C). Blue arrows indicate TLPs. (E) Electron micrograph (low magnification) of RRV-infected *SPCS1* KO cells. ‘V’ indicates the viroplasm. (F) Enlarged view of the area boxed in red in panel (E). Magenta arrows indicate abnormal morphology of TLPs. Scale bars: low magnification, 50 nm; high magnification, 100 nm.

### SPC is important for rotavirus VP7 maturation

VP7 forms the outmost layer of TLPs [24]. It contains an N-terminal signal peptide (**Fig 5A**), which is known to be cleaved between 50 aa and 51 aa [16]. We hypothesized that SPCS1 loss impaired TLP formation by binding to VP7 and interfering with its processing, specifically the cleavage from precursor to mature form. We transfected WT and *SPCS1* KO cells with GFP-tagged VP7 for western blot analysis. NSP4 is another ER-resident protein encoded by rotavirus [25] and we included here as an additional control. Consistent with the TLP formation results, VP7 showed two distinct bands around 65 kDa and 35 kDa. In WT cells, the upper band (precursor form) and lower band (cleaved, mature form) were both present, with the mature form being more abundant (**Fig 5B**). In contrast, in *SPCS1* KO cells, the lower band was nearly absent, and VP7 accumulated predominantly as the upper band, indicating that SPCS1 loss disrupts VP7 processing (**Fig 5B**). To further validate the processing of VP7 was dependent on its cleavage, we introduced a single site mutation (A50V) in the N-terminus of VP7, which was known to prevent the signal peptide removal [16]. As a result, the cleavage mutant impaired the processing of VP7 in WT cells as evidenced by predominant accumulation of the precursor (upper) band, similar to that observed in *SPCS1* KO cells (**Fig 5B**). As expected, the cleavage and processing of WT and A50V mutant VP7 proteins were both impaired in the *SPCS1* KO cells as shown by the nearly undetectable levels of the lower VP7 band (**Fig 5B**), further confirming the essential role of SPCS1 in VP7 processing. To quantify processing efficiency, we also calculated the cleavage ratio efficiency between the WT and A50V mutant VP7 proteins in WT and *SPCS1* KO HEK293T cells by scanning and quantifying the intensities of the bands. In WT cells, the cleavage efficiency of WT VP7 protein reached 84%, while that of A50V mutant VP7 protein was significantly lower at 30.7%. In *SPCS1* KO cells, both forms showed minimal cleavage (**Fig 5C**). Importantly, these results demonstrated that SPC specifically facilitates VP7 processing, and the transition from VP7 precursor to the matured form was dependent on the signal peptide cleavage between residues 50 and 51.

**Fig 5.**
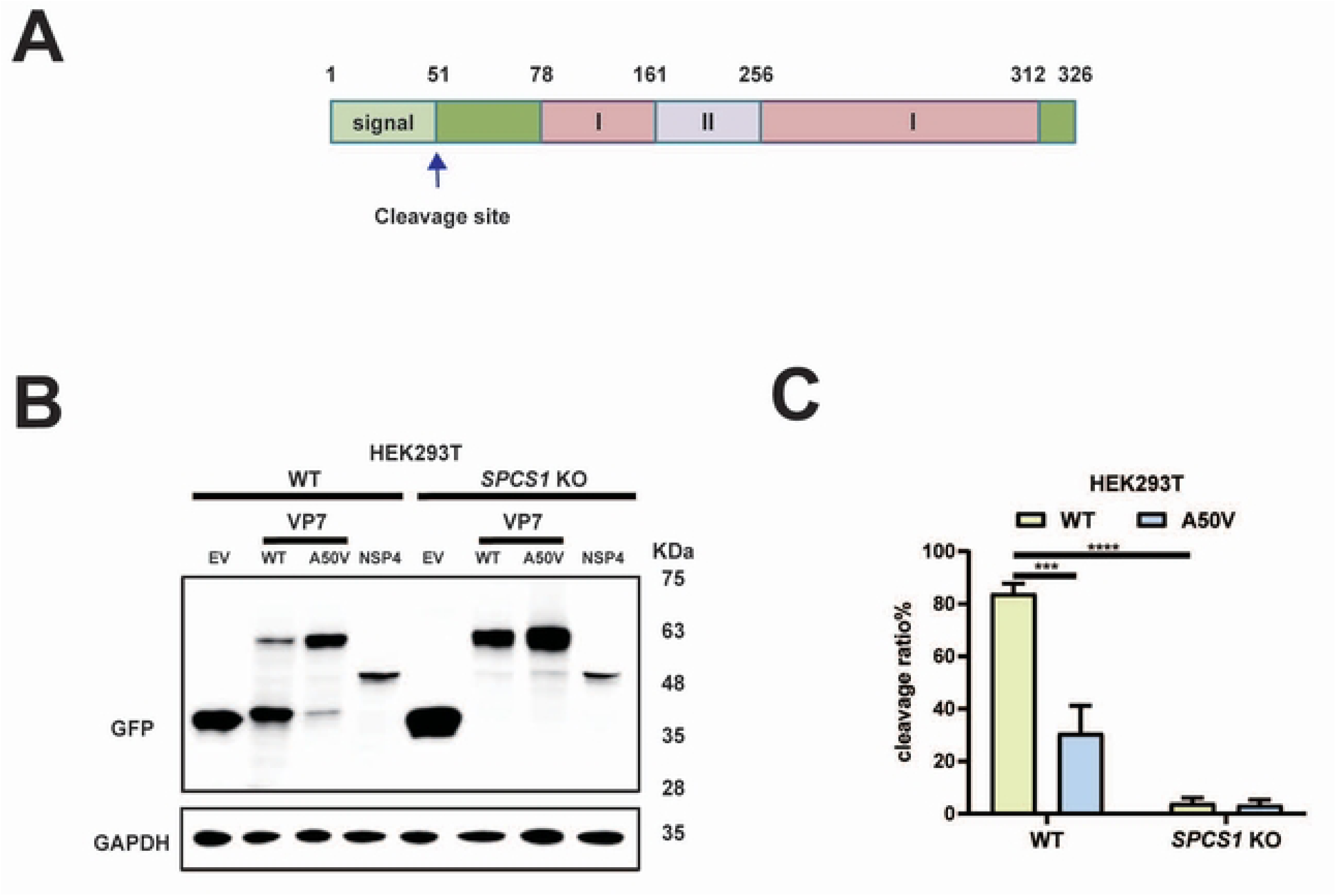
SPC is important for rotavirus VP7 maturation. (A) Schematic diagram of rotavirus VP7, cleavage site was indicated by an arrow. (B) WT and *SPCS1* KO HEK293T cells were transfected with plasmids expressing -EV, - WT VP7, -A50V mutant VP7, -non-structural proteins NSP4, respectively. Those plasmids were all tagged with GFP. At 24 hours post-transfection, cells were harvested for western blot analysis by probing GFP. (C) Quantification of the cleavage ratio of WT and A50V mutant VP7 proteins in WT and *SPCS1* KO HEK293T cells. The results are the averages of data in three independent experiments and plotted as mean ± SD. Statistical significance was determined by two-way ANOVA with Sidak’s multiple comparisons test (***, *P* < 0.001, **** *P* < 0.0001).

### Residue E256 mediates VP7 binding to SPC

To further gain insights into VP7-SPC interaction, we used AlphaFold3 to predict the binding interface of VP7 with SPC complex including SPCS1, SPCS2, SPCS3, and SEC11A (**Fig 6A**). The N-terminus of VP7 looped around SPCS3 and made direct contact with SEC11A, the catalytic subunit (**Fig 6A**). VP7 also had several close-distance interactions with SPCS3 (**Fig 6B**). To experimentally test this prediction, we generated a VP7 mutant with a negatively charged glutamic acid (E) at position 256 changed to a positively charged arginine (R) with site-directed mutagenesis. We transfected WT and E256R mutant VP7 proteins into HEK293 cells and detected the binding efficiency of VP7 and SPC by co-ip assay. The results showed that VP7 specifically co-immunoprecipitated with SPCS1, but not with NSP4, indicating a specific interaction between VP7 and SPC (**Fig 6C**). Notably, compared to WT VP7, E256R mutant had reduced binding to SPCS1, suggesting that residue E256 is crucial for VP7 interaction with SPC. To further investigate whether this interaction is important in the context of rotavirus infection, we leveraged the reverse genetics system and rescued recombinant simian rotavirus SA11 strain that encodes WT and E256R VP7 derived from RRV strain. Recombinant viruses carrying the E256R mutant VP7 both exhibited a 5-fold reduction in infectious titers compared to the parental virus (**Fig 6D**). Importantly, both viruses replicated comparably in *SPCS1* KO cells, suggesting that the E256R substitution attenuates the virus specifically through disruption of the VP7-SPC interaction (**Fig 6D**). Given that single-point mutations are prone to reversion to wild-type sequences, we amplified the *VP7* gene of recombinant parental and VP7 mutant viruses propagated in WT HEK293T cells. A single cycle of replication of the E256R mutant readily gave rise to a mixed population of approximately 33.3% WT sequences (**Fig 6E**). Taken together, these data suggest that the VP7 contact site with SPC is critical for optimal viral growth and potentially explained why we only observed a 5-fold reduction in new progeny.

**Fig 6.**
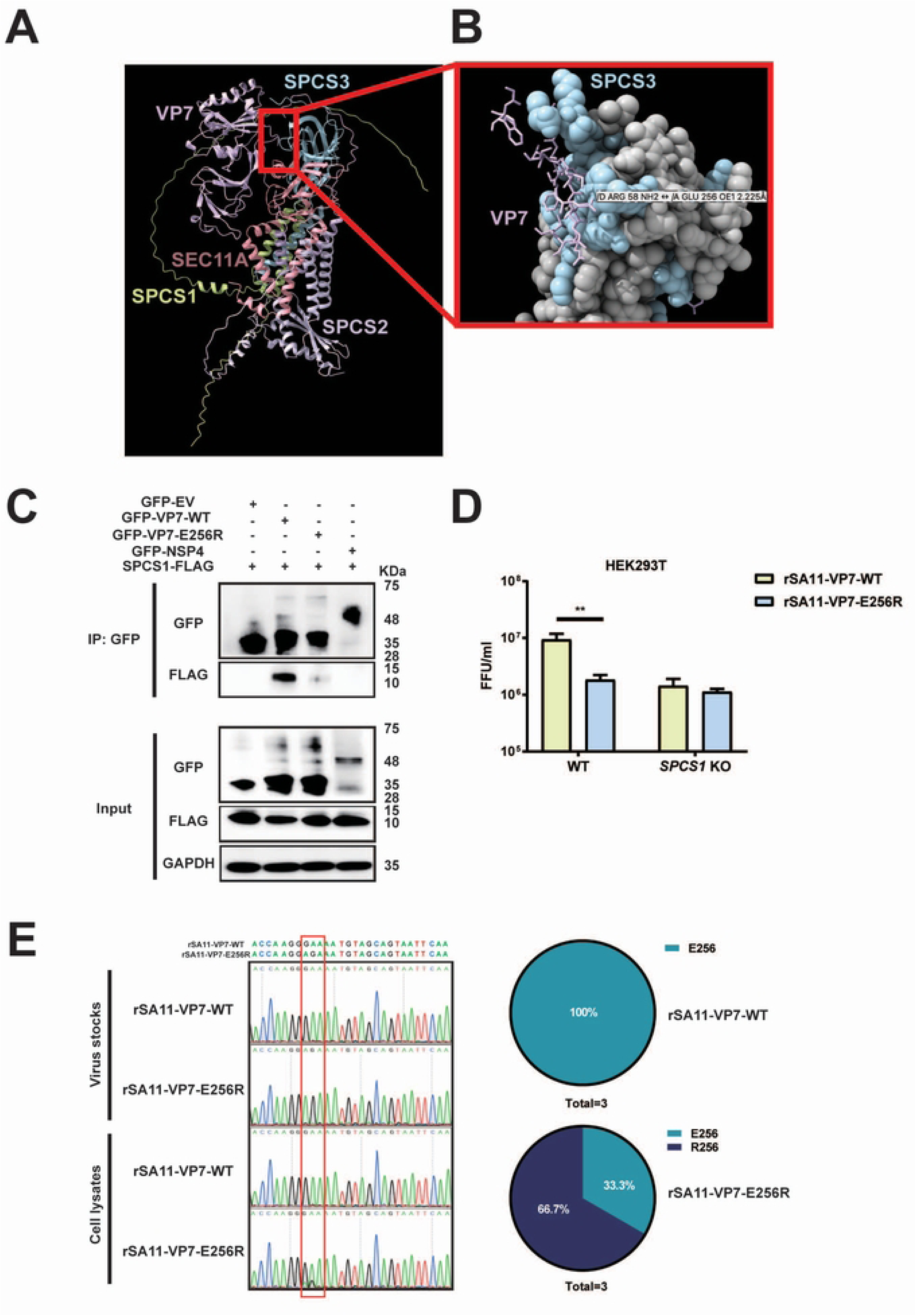
Residue E256 mediates VP7 binding to SPC. (A) Alpha-Fold 3 prediction of VP7 with SPC complex (SPCS1, SPCS2, SPCS3, and SEC11A). (B) The magnification of the interaction site of VP7 and SPCS3 from (A). (C) Co-IP of SPCS1 with VP7 mutants. HEK293 cells were co-transfected with plasmids expressing SPCS1-FLAG with GFP-WT VP7, -E256R mutant VP7, -NSP4 and -EV, respectively. Cell lysates of transfected cells were immunoprecipitated with anti-GFP antibody. The resulting precipitates and whole-cell lysates used for immunoprecipitation were examined by immunoblot using anti-GFP and anti-FLAG antibodies. (D) WT and *SPCS1* KO HEK293T cells were infected with rSA11-WT or rSA11-E256R at an MOI of 3. At 12 hpi, all the cell lysates and supernatants were collected for FFU assay to detect the titers. Statistical significance was determined by two-way ANOVA with Sidak’s multiple comparisons test (***, *P* < 0.001, **** *P* < 0.0001). (E) Samples were collected from WT HEK293T cells infected with rSA11-VP7-WT and rSA11-VP7-E256R in three independent experiments. The purified PCR products targeting the *VP7* gene were subjected to Sanger sequencing, and the results are shown in the chart (right panel). Direct sequencing of *VP7* from rSA11-VP7-WT and rSA11-VP7-E256R viruses served as controls. The red box indicates the codon of mutated amino acids.

## Discussion

SPC plays a very important role in mediating the viral infection. In most studies, it is reported that SPCS1 plays an indispensable role in cleaving signal peptides from viral glycoproteins, facilitating their correct maturation and assembly, especially for flaviviruses [3]. However, the exact role of SPC in rotavirus remains unknown. In this study, we, for the first time, verified that SPCS1 was necessary for rotavirus replication. This study starts with an interactome study of VP7 of a human strain, DS-1, the approach was previously introduced for other viral proteins of RRV [22, 23]. Surprisingly, all the members of SPC family appeared in the interactome, and showed strong interaction with VP7 which was further validated by co-localization analysis and co-ip assay. We validated SPC complex as a whole could mediate RV propagation and also is not strain dependent at least not for the three strains tested. We also demonstrated that SPCS1 deficiency did not affect viral protein or mRNA levels, nor the replication, consistent with previous reports [3, 5].

One recent study showed no virion-like structures of JEV were detected in the ER of *SPCS1* knockout cells, despite comparable levels of viral RNA replication and protein expression [5]. In our study, the outmost layer, VP7, which constitutes TLP, was absent in the viral particles from *SPCS1* KO cells. Unlikely, in RRV-infected *SPCS1* KO cells, instead of seeing no virion-like structures, we observed abundant aberrant viral TLPs, indicating defective TLPs fail to assemble properly, rendering viral particles non-infectious, thereby decreasing the progeny of RV.

Several studies have proven cleavage site of VP7 is between Ala50 and Gln51[19, 26, 27]. A single point mutation that converts Ala50 to Val prevented VP7 processing [16]. However, the host signal peptidase responsible have not been determined. In our study, we define host protein SPC as the mediator of VP7 maturation by processing cleavage of its precursor, as evidenced by the use of an A50V mutant VP7. Rotavirus VP7 is a calcium-dependent glycoprotein that constitutes the outermost layer of the viral capsid and plays a critical role in virus assembly and infectivity. During morphogenesis, VP7 associates with the intermediate VP6 layer and stabilizes the mature triple-layered particle. The proper folding and trimerization of VP7 within the ER are essential for its incorporation onto the DLP, a process that also requires the removal of the ER signal peptide and coordination with the non-structural protein NSP4. In the absence of functional VP7, rotavirus particles remain incomplete and non-infectious, highlighting its indispensable role in virion maturation [16, 17, 27]. Rotavirus NSP4 is an integral ER transmembrane glycoprotein that plays multiple roles in viral morphogenesis. Within the ER, NSP4 functions as a receptor that mediates the budding of double-layered particles from the cytoplasm into the ER lumen, a critical step for the acquisition of the outer capsid proteins VP7 and VP4. NSP4 interacts directly with DLPs, anchoring them to the ER membrane and facilitating their envelopment [28-30]. In addition, NSP4 acts as a viroporin, altering ER membrane permeability and calcium homeostasis, which in turn promotes correct VP7 folding and trimerization [25, 31, 32]. Thus, NSP4 serves as both a viral receptor and a molecular chaperone within the ER, orchestrating the maturation of infectious TLPs [29]. In our study, we found that SPCS1 primarily mediates VP7 rather than NSP4 although both viral proteins reside in the ER by detecting their expressions in *SPCS1* KO cells and interaction with SPCS1. The underlying cause may be that SPC recognizes and cleaves signal peptides based on conserved sequence and structural rules. A typical signal peptide consists of a positively charged N-region, a hydrophobic H-region, and a polar C-region that carries the cleavage site. Cleavage usually follows the (-3, -1) rule, in which small, neutral residues such as Ala, Ser, or Gly occupy the -3 and -1 positions upstream of the scissile bond [1], while bulky or charged residues at these positions strongly reduce processing. VP7 conforms to these rules: its N-terminal signal peptide has an appropriate hydrophobic core and a C-region with favorable (-3, -1) residues, leading SPC to cleave precisely between residues 50 and 51. In contrast, the N-terminal sequence of NSP4 (residues 1–30) behaves as a signal anchor rather than a cleavable signal peptide [25]. Its C-region does not satisfy the (-3, -1) rule, and the hydrophobic segment is longer and more anchor-like, preventing SPC-mediated cleavage. In addition, VP7 interacts with SPC subunits to facilitate recognition, whereas NSP4 shows no detectable interaction (**Fig 6C**), further explaining why SPC processes VP7 but not NSP4.

Although the E256R mutant recombinant virus attenuated viral propagation due to its reduced binding to SPC, the binding is not completely abrogated, indicating there were more sites undefined responsible for the binding, like Arg 58, and complete disruption of SPC-VP7 binding may drastically impair viral propagation. Furthermore, we observed that the recombinant viruses contain E256 mutant VP7 gradually reverted to the wild-type virus during continuous proliferation, indicating the genetic instability of a single-point mutant recombinant virus, the reversion would weaken the phonotype to some degree. Moreover, the binding site prediction was between VP7 and SPC, in addition to the catalytic subunit SEC11A, VP7 had a close interface with SPCS3 at E256 and R58 (**Fig 6A-B**). As a core subunit, SPCS3 participates in cleaving signal peptides from nascent polypeptides. However, SPCS3 is a strong dependency gene across many cell lines, with loss resulting in lethality which was proven by Zhang [3]. Consistently, experimental attempts to generate homozygous SPCS3 knockout clones usually fail. It is evident that the SPC complex functions as an integrated unit to facilitate this process. SPCS1 coordinates the activity of the SPC complex to process VP7, while the exact functional subunit of SPC processing VP7 has not be precisely defined.

Our findings highlight the complexity of the interaction between VP7 and the SPC complex, with evidence suggesting multiple binding sites beyond E256, such as Arg58, contribute to this interaction. Future studies should focus on mapping these additional interaction sites in greater detail to fully understand the molecular interface between VP7 and SPC subunits. In addition, replacing the rhesus RV VP7 with VP7 derived from heterologous rotavirus strains to further reveal the maturation mechanism of rotavirus. Furthermore, the observed genetic instability and reversion of E256 mutant viruses indicate a strong selective pressure to maintain functional VP7-SPC binding, we plan to generate recombinant rotaviruses that lack a portion of the VP7 coding region to prevent the reversion to wild-type parental strains, warranting further investigation into the evolutionary constraints governing this interaction. Given the essential role of SPCS3 as a core SPC subunit and its lethality upon loss, developing conditional knockout or knockdown models could provide valuable insights into its specific contribution to VP7 processing without compromising cell viability. Additionally, dissecting the individual roles of other SPC subunits, including SEC11A and SPCS1, in VP7 cleavage and rotavirus assembly remains critical to fully elucidate the SPC complex’s mechanistic function. Ultimately, such studies could identify novel therapeutic targets within the SPC complex to inhibit rotavirus maturation and propagation.

In conclusion, our study reveals that SPCS1 is required to facilitate VP7 processing to promote rotavirus assembly. For the first time our study provides the evidence that the host factor SPC regulates rotavirus assembly through interactions with viral protein VP7. Our findings provide clues for understanding the molecular mechanism of the assembly of infectious rotavirus and other RNA viruses, offer a novel antiviral strategy, screening for compounds that disrupt SPC-VP7 interaction, especially with the residue 256 of VP7, represents a promising avenue for therapeutic development.

## Materials and methods

### Cells, viruses, and antibodies

Human embryo kidney HEK293T cells (ATCC CRL-3216), were grown in Dulbecco’s modified Eagle’s medium (DMEM) (catalog number 11965118; Gibco) supplemented with 10% heat-inactivated fetal bovine serum (FBS) (catalog number 267820; Avantor). MA104 cells (ATCC CRL-2378) were cultured in Medium 199 (M199, Sigma-Aldrich) supplemented with 10% heat-inactivated FBS, 0.292mg/ml L-glutamine.100 U/ml penicillin, and 100 μg/ml streptomycin at 37°C in 5% CO_2_. Huh7.5 hepatoma cells (JCRB0403) were obtained from the Japanese Collection of Research Bioresources Cell Bank (JCRB), were cultured in DMEM supplemented with 10% heat-inactivated FBS. *SPCS1* KO Huh7.5 cells and *SPCS1* KO HEK293T cells were previously characterized [3] and generously provided by Dr. Michael Diamond at Washington University in St. Louis.

Rotavirus strains including rhesus rotavirus (RRV), simian SA11, and bovine UK were propagated and titrated in MA104 cells [33]. Mouse monoclonal antibody against the rotavirus VP6 was obtained from Santa Cruz (catalog number sc-101363). Rabbit polyclonal antibody against SPCS1 was obtained from proteintech (catalog number 11847-1-AP). Rabbit monoclonal antibody against glyceraldehyde-3-phosphate dehydrogenase (GAPDH) was obtained from cell signaling technology (USA, INC. CA, catalog number 5174T). Rabbit monoclonal antibody against FLAG (catalog number 2368T) was obtained from cell signaling technology. Mouse monoclonal antibody (catalog number sc-101536) against GFP was obtained from Santa Cruz. Alexa Fluor 488-conjugated goat anti-mouse IgG antibodies were obtained from Thermo Fisher (catalog number A32723TR). DAPI was obtained from Thermo Fisher (catalog number 62248).

### Plasmids

Plasmids expressing of rotavirus proteins (NSP4, DS1-VP7) with GFP tags, pG-LAP6-RRV-GFP were described previously [34, 35]. pG-LAP6-RRV-VP7, pG-LAP6-mutant VP7 at 50aa (A mutated into V) were synthesized by GenScript Biotech company (NJ, USA). The set of plasmids for RV rescue: pT7-SA11 of 11 genes and C3P3-G3 were prepared as described previously [36]. Mutant pG-LAP6-RRV-VP7(E256R), pT7-RRV-VP7(E256R) plasmids were generated using a Quikchange II site directed mutagenesis kit (Agilent). The plasmid expressing C-terminally Flag-tagged SPCS1 was generously provided by Dr. Michael Diamond at Washington University in St. Louis. All the plasmids are purified by using a QIAGEN Plasmid Maxiprep kit per the manufacturer’s instructions.

### Inoculation of rotavirus and treatment

WT or *SPCS1* KO HEK293T cells cultured in 24-well plate (1 × 10^5^ cells /well) in DMEM complemented with 10% (vol/vol) FCS and 100 IU/ml penicillin-streptomycin for 24 h to reach 90% confluency. Rotavirus (1 × 10^8^ FFU/ml) was activated with 5 μg/ml of trypsin at 37°C for 20 minutes. The cells were infected with rotavirus at an MOI of 3, incubation at 37°C for 1 h and followed by three times washing with sterile PBS to remove free virus particles. Subsequently, serum-free DMEM medium were added to the infected cells and incubated for further 12 hours.

### DNA transfection

WT and *SPCS1* KO HEK293T cells (90% confluent) were transfected with plasmid DNA by using Lipofectamine™3000 Transfection reagent (Thermo Fisher, catalog number L3000075) according to the manufacturer’s instructions. Viral infection was performed at least 24 h post transfection.

### siRNA transfection

WT cells or *SPCS1* KO HEK293T cells (60% confluent) were transfected with ON-TARGET plus siRNA pool targeting *SPCS2*, *SEC11A*, *SEC11C* (available in Dharmacon LQ-005932, LQ-020897, LQ-006038, respectively) using Lipofectamine™ RNAiMAX Transfection reagent according to the instruction (Thermo Fisher, Catalog number 13778150). 72 hours later, the transfection mixture was removed and the cells were washed twice with MEM and infected with rotavirus.

### Reverse transcription and quantitative PCR (RT qPCR)

Intracellular rotavirus RNA was isolated by using a RNeasy RNA isolation kit (Qiagen, Valencia, CA). cDNA Reverse use High-Capacity cDNA Reverse Transcription Kit (Catalog number 4368814). To quantitate the level of rotavirus RNA, a real-time RT-PCR assay was performed by using a TaqMan Fast Advanced Master Mix for qPCR targeting NSP5 (Thermo Fisher, Catalog number 4444963) as previously described[22, 34] and SYBR Fast Advanced Master Mix (Thermo Fisher, Catalog number 4385612) for qPCR targeting SPCS2, SEC11A, SEC11C. Primers were listed in **Table 2**.

**Table 2.**
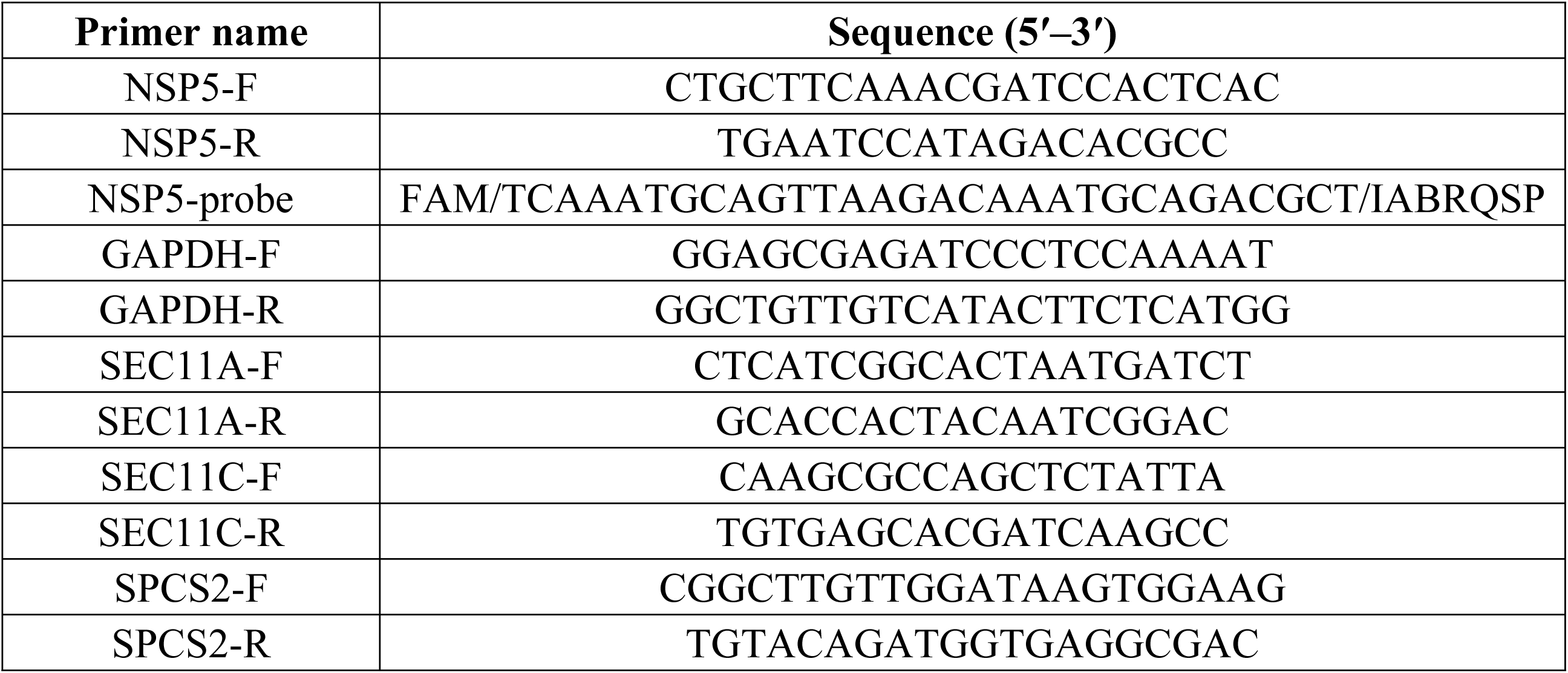
Primer sequences used in this study. This table lists the sequences of primers and probe used for qPCR analysis in the study. Primer names are listed along with their corresponding sequences in the 5’ to 3’ direction. The NSP5 probe is labeled with a fluorescent dye FAM and quencher IABRQSP for qPCR detection using a TaqMan probe, while the remaining primers targeting *GAPDH*, *SEC11A*, *SEC11C*, *SPCS2*, are used for SYBR Green qPCR detection.

### Virus titration by focus forming unit assays

The positive cells were enumerated and virus titers were expressed as focus forming unit (FFU) per ml as previously described [22]. Activate virus with trypsin (5 μg/ml) at 37°C for 15 - 20 min. Activated virus were serially diluted 10-fold in M199 serum-free medium and added to confluent monolayers of MA104 cells seeded in 96-well plates for 1 h at 37 °C. Virus inoculates were discarded and replaced with M199 serum-free medium, then incubated for 18 h. Rotavirus-infected MA104 cells were then fixed with 10% paraformaldehyde for 30 min at room temperature and permeabilized with 1% triton-X for 3 min at room temperature. After washing three times with PBS, the cells were incubated with mouse anti-VP6 monoclonal antibody (1:2000) and HRP-conjugated goat anti-mouse IgG secondary antibody (cell signaling, MA, USA) (1:1000). Viral antigenic foci were visualized with the AEC substrate kit (Vector laboratories, California, USA, cat number SK-4205) according to the manufacture’s instruction. The positive cells were enumerated and virus titers were expressed as FFU per ml.

### Western blot analysis

Cells were lysed by lysis buffer contain 1% protein inhibitor (Thermo Scientific, MA, USA, cat number J61852-XF) 30 μg cell lysate protein of each sample was subjected to 10% sodium dodecyl sulfate–polyacrylamide gel electrophoresis (SDS-PAGE) and transferred onto 0.45-μm nitrocellulose membranes (Bio Rad, CA, USA). Then, the membranes were incubated with 5% BSA at room temperature for 2 h, then washed by Tris-buffered Saline with Tween 20 (TBST) for three times, incubated with correspondent primary antibodies at 4°C overnight, followed by incubation with horseradish peroxidase-conjugated goat anti-mouse (or rabbit) IgG secondary antibody at room temperature for 2 h. Detection was performed using the ECL System (Thermo Fisher Scientific). followed by imaging with an Chemi Doc MP imaging system (Bio Rad, CA, USA).

### Coimmunoprecipitation assays

HEK293 cells were transfected with target plasmids and harvested for further 24 h after transfection. Transfected cells were washed twice with phosphate-buffered saline (PBS) and scraped into lysis buffer (NP40) containing 1 X protease inhibitor (Thermo Fisher MA, USA). After incubation for 30 min on ice, the lysate was centrifuged to remove the cell debris. Dynabeads protein G (Thermo Fisher, MA, USA, Catalog number 10003D) 10 μl per sample was prepared according to the instruction of the manufacturer and added with 2 μg of anti-GFP antibody (Santa Cruz, CA, USA) at 4°C diluted in 200 μl PBS with Tween 20, incubate with rotation for overnight at 4 °C. Equal amounts of lysates were used for the Co-IP assay by incubating overnight with Dynabeads protein G beads coupled with GFP antibody at 4°C. Wash Dynabeads with cold-PBS twice. The immune complexes were denatured by 37°C, 20 mins. Samples were then subjected 4-20% SDS-PAGE and analyzed by western blot.

### Immunofluorescence staining

Immunofluorescence staining of viroplasm in infected cells was performed by immunofluorescence labeling staining using an in-house anti-NSP5 antibody [37, 38]. Briefly, the infected WT and *SPCS1* KO HEK293T cells were fixed with 4% formaldehyde for 30 min at room temperature and discard washed by PBS for two times. Add 1% triton-X and incubate for 5 min at room temperature to permeabilize cells. After being washed three times in PBS, then incubation with guinea pig anti-NSP5 polyclonal antibody (1:1000) for an hour at 37 ◦C. After being washed three times in PBS, the cells were reacted with 488-conjugated goat anti-guinea pig IgG secondary antibody (Thermo Fisher, cat number A18769) (1:1000) for 1 h at 37 ◦C and then incubated with DAPI (1:1000) for 5 min. Visualize staining of tissue, acquisitive and analysis image were conducted by a fluorescence microscope. Co-localization was analyzed by Volocity 7.

### Transmission electron microscopy

WT or *SPCS1* KO HEK293T cells cultured in 6-well plate (5 × 10^5^ cells /well) and infected with RRV at an MOI of 3. At 12 hours post-infection (hpi), adherent cells were trypsinized and collected by centrifugation. Then the infected cells were fixed in 2% paraformaldehyde/2.5% glutaraldehyde (Ted Pella Inc., Redding, CA) in 100 mM cacodylate buffer, pH 7.2 for 2 h at room temperature and then overnight at 4°C. Samples were washed in cacodylate buffer and postfixed in 1% osmium tetroxide (Ted Pella Inc.)/ 1.5% potassium ferricyanide (Sigma, St. Louis, MO) for 1 h. Samples were then rinsed extensively in dH_2_O prior to en bloc staining with 1% aqueous uranyl acetate (Ted Pella Inc.) for 1 h. Following several rinses in dH_2_O, samples were dehydrated in a graded series of ethanol and embedded in Eponate 12 resin (Ted Pella Inc.). Ultrathin sections of 95 nm were cut with a Leica Ultracut UCT ultramicrotome (Leica Microsystems Inc., Bannockburn, IL), stained with uranyl acetate and lead citrate, and viewed on a JEOL 1200 EX transmission electron microscope (JEOL USA Inc., Peabody, MA) equipped with an AMT 8-megapixel digital camera and AMT Image Capture Engine V602 software (Advanced Microscopy Techniques, Woburn, MA).

### Generation of recombinant rotaviruses

Recombinant rotaviruses were generated as described [36, 39]. Briefly, BHK-T7 cells were seeded in 12-well plates 48 h, and then were co-transfected with 0.4 μg of either RV rescue pT7 plasmid, except pT7-NSP2 and pT7-NSP5, which were added at 1.2 μg, and 0.8 μg of the plasmid pCMVScript-NP868R-(G4S)4-T7RNAP (C3P3-G3) with using 14 μl of TransIT-LT1 (Mirus Bio LLC) transfection reagent per reaction. All the plasmids and transfection reagents were mixed and then incubated at room temperature for 20 min. After that, the transfection mixture was added into medium of BHK-T7 monolayers. Sixteen hours later, after two washes with FBS-free medium, 800 μl of serum-free DMEM was added to the transfected-BHK-T7 cells. Twenty-four hours later, 5×10^4^ MA104 in 200 μl of serum-free DMEM were added to the well, along with 0.5 μl/ml of porcine pancreatic type IX-S trypsin (Sigma-Aldrich). MA104 and BHK-T7 cells were cocultured for 72 h, after which they were frozen and thawed three times. After removal of cell debris, the lysate was activated with 2.5 μg/ml of trypsin to infect a 3-day-old monolayer of MA104 cells, supplemented with 0.5 μg/ml of trypsin. MA104 cells were incubated at 37°C for 5 days or until cytopathic effects were observed.

### Statistical analysis

Statistical analysis was performed using GraphPad Prism version 7.0.2 (Software, San Diego, CA, USA) to determine the significance in differences. Comparisons between groups were performed using unpaired two-way ANOVA with Sidak’s multiple comparisons test or one-way analysis of variance. *P* < 0.05 represents a statistically significant difference. All data are expressed as the mean ± standard deviation (SD).

## Acknowledgments

This study is supported by the National Institutes of Health (NIH) grant R01 AI188179 and Gates Foundation grant INV-075616 to S.D. We thank Dr. Michael Diamond at Washington University who provided *SPCS1* KO HEK293T and *SPCS1* KO Huh7.5 cells.

## Author contributions

Conceptualization: X.Z. L.S.T., and S.D. Methodology: X.Z., L.S.T., and W.B. Investigation: X.Z. and L.S.T. Visualization: X.Z. Supervision: S.D. and B.L. Writing-original draft: X.Z. Writing-review and editing: X.Z., S.D.

---

**S1 Fig. Localization of host protein and viral protein *in vitro*.**

Plasmids expressing -VP7 and REEP5, SEC11C, SPCS2, -VP3 and PFDN4, -NSP1 and SAMD9, -VP6 and ECE1 were co-transfected into HEK293 cells, respectively, and subjected to IFA detection. Viral proteins were tagged with GFP, shown as green fluorescence, while host proteins were tagged with RFP, shown as red fluorescence. Nuclei were counterstained with DAPI. Scale bar, 170 μm.

**S2 Fig. Knockdown efficiency of SPC components in WT and *SPCS1* KO HEK293T cells.**

WT and *SPCS1* KO HEK293T cells were transfected with siRNAs targeting against *SPCS2*, *SEC11A*, *SEC11C* and a scrambled siRNA at the concentration of 20 nM. At 72 hours post-transfection, cells were collected for RNA extraction, and relative mRNA expression levels were measured by qPCR. The relative expression was normalized to GAPDH. Results are the average of data from two independent experiments and plotted as mean ± SD. Statistical significance was determined by two-way ANOVA with Sidak’s multiple comparisons test (*, *P*< 0.05, **, *P* < 0.01; ***, *P* < 0.001, ****, *P* < 0.0001).

**S3 Fig. RNA and protein level in wildtype and *SPCS1* KO Huh7.5 cells during rotavirus infection.**

(A) WT and *SPCS1* KO Huh7.5 cells were infected with RRV at an MOI of 3. At 4, 8, 12 hpi, the cells were collected for qRT-PCR analysis of viral mRNA level by detecting NSP5. NSP5 level was normalized to GAPDH. The result was representative of one independent experiment. (B) WT and *SPCS1* KO Huh7.5 cells were infected with RRV at an MOI of 3. At 4, 8, 12 hpi, the infected and mock cells were harvested for western blot analysis of viral protein levels by detecting VP6.

**S4 Fig. Transmission electron microscopy images of viral particle morphology.**

Transmission electron micrographs of RRV particles in HEK293T cells. (A) Single-layered particles indicated by blue arrows surrounding the viroplasm. ‘V’ indicates the viroplasm. (B) DLP, indicated by blue arrow. (C) TLPs, indicated by blue arrow. (D) The budding process of DLP morphing into TLP, indicated by blue arrow, ‘V’ indicates the viroplasm. Scale bar, 100 nm.

**Dataset 1. Raw data of LC-MS/MS for DS-1 VP7**

Peptide-spectrum matches were identified against the NCBI human protein database, the raw data were listed in three columns. The Spectra table lists individual peptides with their observed and theoretical precursor masses, charge states, modifications. The Proteins table summarizes all identified proteins, including cumulative log-probabilities, best scores, total spectral counts, sequence coverage, and numbers of unique and modified peptides. The Summary sheet includes information on the data file, search parameters. These raw data collectively describe the peptide and protein composition detected in the analyzed sample.

## Notes

### Competing Interest Statement

The authors have declared no competing interest.

